# Structural analysis of pathogenic missense mutations in *GABRA2* and identification of a novel *de novo* variant in the desensitization gate

**DOI:** 10.1101/678219

**Authors:** Alba Sanchis-Juan, Marcia A Hasenahuer, James A Baker, Amy McTague, Katy Barwick, Manju A Kurian, Sofia T Duarte, NIHR BioResource, Janet Thornton, F Lucy Raymond

**Author notes:** These authors contributed equally to this work (Alba Sanchis-Juan, Marcia A Hasenahuer). **Corresponding author**: Professor F Lucy Raymond.

## Abstract

Cys-loop receptors are vital for controlling neuronal excitability in the brain and their dysfunction results in numerous neurological disorders. Recently, six *de novo* missense variants in *GABRA2* gene, a member of this family, have been associated with early infantile epileptic encephalopathy (EIEE) and intellectual disability with seizures. Here, using whole-genome sequencing we identified a *de novo* missense variant in *GABRA2* gene in a patient with EIEE and developmental delay. We perform protein structural analysis of the seven variants and show that all the mutations are in the transmembrane domain, either close to the desensitization gate, the activation gate or in inter-subunit interfaces. Further investigations demonstrated that the majority of pathogenic variants reported are at equivalent positions in other Cys-loop receptors, emphasizing the importance of these residues for the adequate function of the receptor. Also, a comparison of the distribution of the mutations in all the Cys-loop receptors showed that pathogenic variants are more common in the transmembrane helices, more specifically in the M2 helix, highlighting the importance of this segment. Our study expands the clinical spectrum of individuals with pathogenic missense mutations in *GABRA2*, defines the regions where pathogenic mutations are in the protein structure, and highlights the value of considering sequence, evolutionary, and structural information from other Cys-loop receptors as a strategy for variant interpretation of novel missense mutations in *GABRA2*.

## Introduction

The dynamic partnership between excitatory principal cells and inhibitory interneurons is essential for proper brain function and needs to be maintained to avoid pathological consequences. Cys-loop ligand-gated ion-channels are receptors activated by neurotransmitters and play an important role in the development and activity of the central and peripheral nervous systems (1). These receptors mediate excitatory and inhibitory synaptic transmissions depending on the distribution of ions at either side of the membrane and the membrane potential of the cell. There are two broad classifications of Cys-loop receptors, each with different electrophysiological properties: firstly cation-selective receptors, corresponding to nicotinic acetylcholine, serotonin (5-HT_3_) and zinc-activated receptors and secondly anion-selective members, that include glycine (Gly) receptor and γ-aminobutyric acid receptors type A (GABA_A_) and C (GABA_C_) (1, 2).

Cys-loop receptors are pentameric channels formed by five chains, each comprising an extracellular domain (where the ligand binds) and a transmembrane (TM) domain consisting of four TM helices (TMH), M1 to M4 (1). The pentamers can be assembled from different gene products that encode different subunits, but all belong to the same type of receptors. For example, for GABA_A_ receptors (GABA_A_R), there are 19 genes encoding different subunits (α1-α6, β1-β3, γ1-γ3, ρ1-ρ3, δ, ε, π, θ) (3, 4), and the most abundant complex in adult human receptors is a combination of two α, two β and one γ subunit (5). Cys-loop receptors are conserved in humans and across species (6, 7). Dysfunctions of these receptors demonstrate a critical role in neurological development. Pathogenic variants in genes encoding receptors of nACh (*CHRNA2, CHRNA4* and *CHRNB2*) and GABA_A_ (*GABRA1, GABRB1, GABRB2, GABRB3, GABRG2*, and *GABBR2*) have previously been associated with human epilepsy (8), and mutations in Gly receptors (*GLRA1* and *GLRB*) cause hyperekplexia (9, 10).

For many years, *GABRA2*, that encodes for GABA_A_R subunit α2 (GABA_A_R α2), remained as a candidate gene for epilepsy. Multiple genome-wide association studies identified non-coding single nucleotide polymorphisms in *GABRA2* associated with increased risk for epilepsy (11, 12), as well as alcohol dependence and brain oscillations (13, 14). Also, decreased expression of *GABRA2* in *Scn1a* +/- mice served as a model of Dravet syndrome (15, 16), and increased expression observed in Pumilo-2 deficient mice resulted in enhanced seizure susceptibility and the manifestation of epilepsy in the hippocampus (15). *GABRA2* is highly expressed in the hippocampus, especially in early development, and is localized to the cell soma to mediate synaptic transmission (17, 18). Recently, seven individuals with missense variants in *GABRA2* and Early Infantile Epileptic Encephalopathy (EIEE) have been reported separately (19–21). Six variants were reported in total: five were *de novo* and one was present in two affected siblings, inherited from their mosaic father.

Here, we describe the eighth individual with EIEE and a novel *de novo* missense variant in *GABRA2* identified by whole-genome sequencing (WGS). We perform protein structural mapping and analysis of all the seven variants in the GABA_A_R and investigate the effect of the novel variant in the protein. We analyze the presence of variants in equivalent positions in other members of the Cys-loop receptors family and compare their reported effect. Furthermore, because the seven variants cluster in the TM domain, we also analyze the distribution of previously reported pathogenic variants of the Cys-loop receptors. Our results demonstrate the utility of performing variant interpretation by gathering together sequence, evolutionary, and structural information from homologous Cys-loop receptors to facilitate the characterization of novel candidate missense variants.

## Results

### Clinical evaluation

A female was born at term (41 weeks) by Caesarian section. There was no family history of disease and no consanguinity reported. At delivery, Apgar scores were 9 and 10 and her head circumference was 34 cm (25^th^-50^th^ percentile). She presented at 15 months of age with non-specific EIEE and global hypotonia. Seizures occurred during sleep, with and without fever, and were tonic with upward eye deviation, or eye and head deviation to either side. MRI performed at 19 months was normal. EEG at onset of epilepsy was normal, but later investigations showed slow and irregular background activity, without paroxysmal activity. Developmental delay emerged, especially affecting language (comprehension and expression), but also behavior disturbance including hyperactivity, repetitive routines or rituals and marked disturbance with changes in the environment. She also developed hand stereotypies (waving, finger repetitive movements). She was treated with sodium valproate and clobazam that controlled her seizures. Currently, she is 10 years old and has severe impairment of language, hand stereotypies, disruptive behavior and repetitive movements. All routine genetic analyses were negative. Additional clinical details for this individual are compared to the previously reported cases (21) in Supp. Table 1.

### Identification of a novel *de novo* mutation in *GABRA2*

A novel *de novo* missense mutation was identified in *GABRA2* gene at genomic position Chr4:46305494 G>A, NM_000807.2:c.839C>T, NP_000798.2:p.Pro280Leu by whole genome sequencing trio analysis of DNA from both parents and the child. The variant was confirmed by Sanger sequencing (Supp. Figure S1). No other candidate variant was identified in this individual from trio analysis. The gene was observed to be constrained for loss-of-function (LOF) variation (with pLI = 1 and observed/expected score = 0.05) and missense variation (Z score = 3.13) in gnomAD (22), especially in the TM region and ligand binding domain of the neurotransmitter-gated ion-channel (Figure 1A) (23).

Pro280 is located in M2, the second helix in the TM domain (Figure 1B,C). It is highly conserved across species (Figure 1B) and across different receptors from the GABA family in human (Supp. Figure S3A). This variant is absent in gnomAD database (Figure 1A) and was predicted to be damaging by CADD, Sift and Polyphen-2 (with scores of 29.4, 0.02 and 1 respectively).

**Figure 1.**
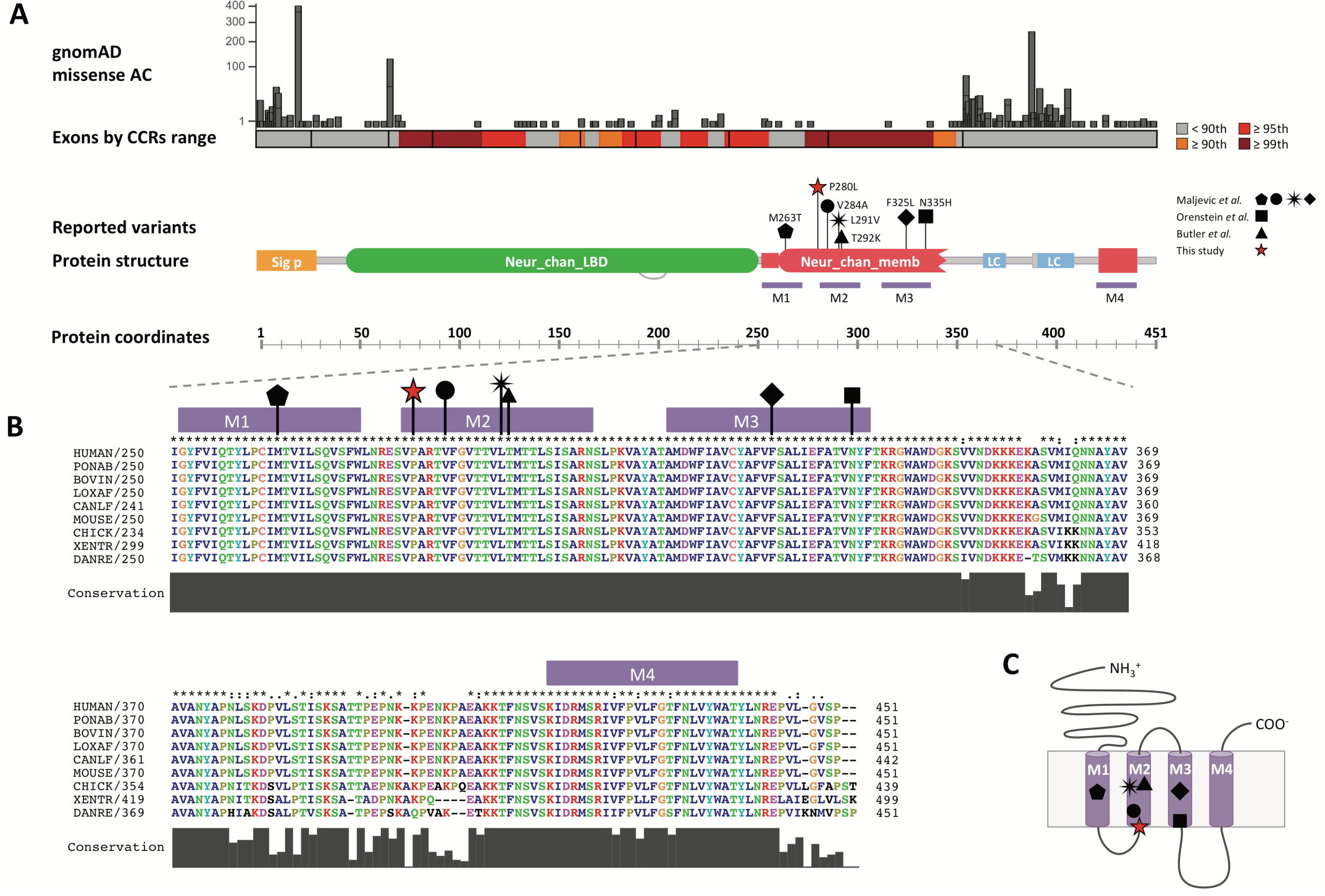
(A) Schematic diagram showing the domains architecture in GABA_A_ α2. At the top, allele count of missense variants obtained from gnomAD are represented. The six *missense* variants in the seven individuals reported in the literature and the novel *de novo* one are indicated with different symbols (triangle, square, circle, diamond, pentagon, black star and red star), and cluster in the TM region, which is highly constrained for variation in control population (see also Supp. Figure 2A). Coordinates of the protein domains were obtained from Pfam (https://pfam.xfam.org). Sig_p = signal peptide; Neur_chan_LBD = neurotransmitter-gated ion-channel ligand binding domain; Neur_chan_memb = neurotransmitter-gated ion-channel TM region; LC = low complexity; TM = transmembrane. **(B)Alignment of the TM region of GABA**_**A**_ **α2 subunit across representative vertebrate species.** The region where the three reported *de novo* variants are is evolutionary conserved through these species. Colors are by amino acid properties. Ponab = orangutan; Bovine = cow; Loxav = elephant; Canlf = dog; Chick = chicken; Xentr = xenopus; Danre = zebrafish. **(C) Schematic representation of GABA**_**A**_ **α2 TMH.** The locations of the seven missense variants are approximate.

### Structural characterization of the novel and previously reported variants in GABA_A_R α2

Protein structural mapping of the seven variants revealed their proximity to the desensitization gate, the activation gate or the M1-M3 inter-subunit interfaces of the receptor.

The variants near the desensitization gate are Pro280Leu, Val284Ala and Asn335His. This gate is defined by the region between −2’ and 2’ positions of the receptor and modulates the conductance and ion-selectivity, as observed in the structures of GABA_A_ and Gly receptors (24–26). Pro280 is located at the TM region of the receptor, in the N-terminal end of the M2 helix in the cytoplasmic facing leaflet (Figure 2), and corresponds to the specific −2’ position. The side chains of the amino acids in this position (Pro in the case of α1-α6 and γ1-γ3, Ala in β1-β3), one coming from each chain of the pentamer, altogether define the inner narrowest constriction of the pore, known as the −2’ ring (26), as shown in Figure 2B and C. A reduction of the diameter of the pore is observed upon the introduction of this variant (Supplemental Figure 4A,B). Small non-polar amino acids (proline and alanine) are conserved at the −2’ position in human GABA_A_ and Gly anion-selective receptors (Supp. Figure 3A), suggesting the necessity of maintaining the size, shape and absence of charge in the inner constriction of this pore for the adequate transit of anions. In contrast, in the cation-selective receptors, a negatively charged amino acid (glutamate) with a longer and more flexible side chain is usually observed in position -1’ (Supp. Figure S3A). Val284, corresponds to the precise 2’ position. In this case, the side chains of the amino acids in this position do not define a ring, but establish inter-subunit Van der Waals (VDW) interactions with M2 helices from the β3 subunits (Figure 2C) (26). Having an alanine instead of a valine at this position affects the radius of the pore differently depending on the arrangement of the different subunits and the state of the receptor (Supplemental Figure 4A,C). Asn335 is located at the C-terminus of the M3 helix, on the inner side of the TM region, at the base of the helical bundle. This amino acid participates in inter and intra-subunit interactions with residues in the M2-M3 loops of β3 and γ2 chains. Asn335 also presents VDW interactions with Val279, right before the −2’ residue in M2 (Pro280) (Figure 2C).

**Figure 2.**
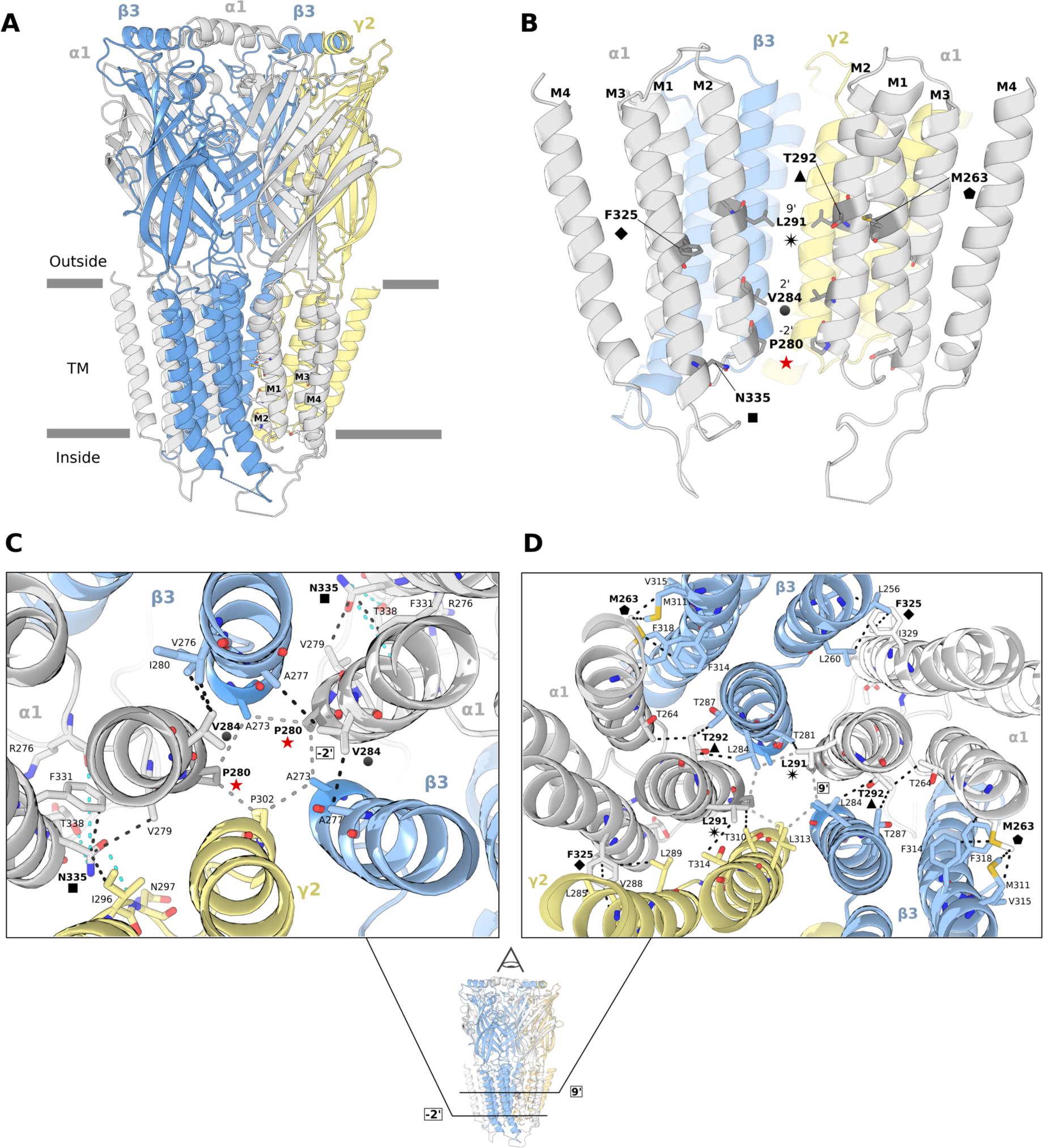
(A) The GABA_A_ α1β3γ2 receptor in closed state. M1 to M4 helices are depicted over the homolog human GABA_A_R α1 chain (PDB ID 6HUG). The two α1 chains are in gray, the two human β3 chains in blue and the human γ2 in yellow, with the approximate boundaries of the TM in gray bars. **(B) Side view of the transmembrane helices of the GABA**_**A**_ **α1β3γ2 receptor in closed state.** The structural location of the variants is depicted consistently using the same symbols as in Figure 1A. For a clearer view, the chain between two α1 subunits is hidden. **(C) Down the pore axis view of the variants nearby the −2’ ring** (Pro280Leu, Val284Ala, Asn335His), and **(D) the 9’ ring** (Met263Thr, Leu291Val, Thr291Lys, Phe325Leu). VDW interactions for the residues where the variants fall are in black dashed lines (except for the rings, that are in gray) and hydrogen bonds are in cyan.

The variants near the activation gate included Leu291Val and Thr292Lys. This gate has been defined as the ring formed by the leucines at the 9’ position, which is important for the transition between opened and closed states of the ion-channel (27). The precise 9’ position corresponds with Leu291, and is located in the pore lining M2 helix of the TM domain (Figure 2B,D). Having a valine instead of a leucine at this residue slightly increases the diameter of the pore (Supp. Figure 4A,D), which would alter the VDW interactions that define the 9’ ring as well as the inter-subunit interactions with the threonines from the neighboring β3 and γ2 chains. Thr292Lys, which is adjacent to Leu291, is involved in inter and intra-subunit VDW interactions with Thr264 in M1 of its same α1 subunit and the β3 subunit residues Leu287 and Leu284, being the last one part of the 9’ ring (Figure 2B,D).

The variants in M1-M3 inter-subunit interfaces were Phe325Leu and Met263Thr. Met263 is located at the middle of M1 helix, approximately at the same height of the 9’ activation gate of M2. It is involved in a network of VDW interactions with residues from the M3 helix of the neighboring β3 subunit. It is near the Thr264, which interacts with the Thr292 (where the mutation Thr292Lys is) (Figure 2B,D). Met263 is also part of the recently reported low affinity binding site for benzodiazepines, a group of sedative anticonvulsant drugs (28–30). In the desensitized GABAA α1β3γ2 receptor bound to Diazepam (DZP), Met263 is part of the opening of the pocket to the external molecular surface and its lateral chain sulfur establishes direct interaction with the diazepine ring of DZP (Supp. Figure 5). Having a smaller threonine with a polar uncharged lateral chain instead of methionine would affect the size and shape of the entrance to this pocket and the interactions established with the benzodiazepines. It can also be observed, deeper in the pocket, that the benzene ring of DZP also interacts with Thr292. Lastly, Phe325, is at the middle of M3 helix, also approximately at the same height of the 9’ activation gate of M2. It is also involved in a network of VDW interactions with residues of M1 from either β3 or γ2 (Figure 2B,D). Some evidence points to the α1 residues Ser297 and Ala318 in the α1-M3 β3-M2 TM interface as critical for the receptor modulation by diverse anesthetics molecules (31).

### The novel variant Pro280Leu affects the diameter of the channel pore in the desensitization gate

We introduced the mutation Pro280Leu in the cryo-EM structure of GABA_A_R α1β3γ2 closed and desensitized structures (Figure 3 and Supp. Figure 4). We considered four situations: i) where α1 is not mutated (α1_P280_), ii) where the modified α1 subunit is between β3 and γ2 (α1^β3γ2^_P280L_), iii) where both α1 subunits are modified (α1^β3γ2-β3β3^_P280L_), and iv) where the mutated α1 subunit is between β3 and β3 (α1^β3β3^_P280L_). In Figure 3, only situations i-iii in the closed state of the receptor are shown for simplicity.

**Figure 3.**
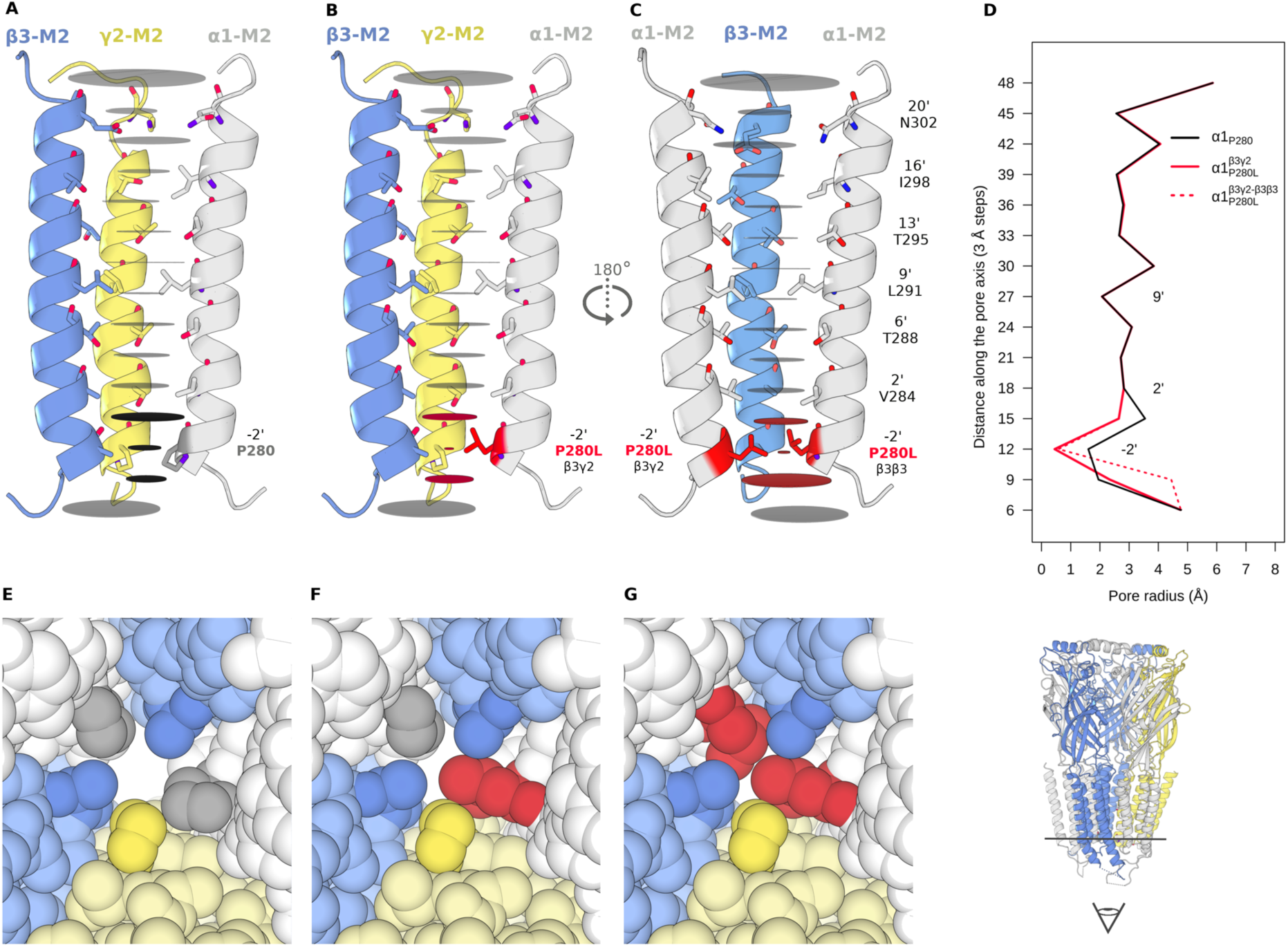
Radii profile through the TM pore axis by 3 Å steps. Representation of M2 pore lining residues shown in sticks for: **(A)** non-mutant GABA_A_ α1β3γ2, **(B)** mutant α1 subunit is between β3γ2 (α1^β3γ2^_P280L_) and **(C)** both α1 subunits are mutated (α1^β3γ2-β3β3^_P280L_). The pore radii at each step along the pore vertical axis is represented with horizontal gray discs. For clarity, only three M2 helices are represented in each situation. The steps with affected radii upon Pro280Leu are colored in red in (B) and (C). **(D)** Pore radius plotted as a function of longitudinal distance along the pore axis, vertically aligned to match the steps in (A), (B) and panels. The biggest radii reductions can be observed at 12 Å (−2’ position) from 1.60 Å in α1_P280_ to: 0.58 Å (Δ=1.02 Å) in α1^β3γ2-β3β3^_P280L_ and 0.45 Å (Δ=1.15 Å) in α1^β3γ2^_P280L_. Also, reductions can be observed at 15 Å (region between −2’ and 2’ positions) from 3.55 Å in α1_P280_ to: 2.64 Å (Δ= 0.91Å) in α1^β3γ2-β3β3^_P280L_ and 2.64 Å (Δ= 0.91Å) in α1^β3γ2^_P280L_. Increments in the radii are observed at 9 Å from 1.95 Å in α1_P280_ to: 4.44 Å (Δ=2.49 Å) in α1^β3γ2-β3β3^_P280L_ and 2.33 Å (Δ=0.38 Å) in in α1^β3γ2^_P280L_. This step at 9 Å corresponds to the vestibule region adjacent to the −2’ ring in the inner cytoplasmic side. Zoom over the inner cytoplasmic vestibule of the pore defined by the amino acids of the −2’ ring for **(E)** non-mutant GABA_A_ α1β3γ2, **(F)** mutant α1 subunit is between β3γ2 (α1^β3γ2^_P280L_) and **(G)** both α1 subunits are mutated (α1^β3γ2-β3β3^_P280L_). The color for each subunit chain matches the ones used in Figure 2.

The larger non-polar amino-acid chain of the leucine narrowed the constriction region compared to the non-mutated receptors (Figure 3D and Supp. Figure 4A,B) measuring the radius at 3 Å steps along the pore axis. The largest changes in all the situations occurred at 12 Å (−2’) and 15Å. For the desensitized structure of the receptor that has GABA and DZP, this reduction can also be observed at 18Å (2’).

FoldX stability calculations showed higher ΔΔG values, which correspond with higher destabilizing effects, in the desensitized form compared to the closed one (Supp. Table 3). These showed a tendency of slightly higher ΔΔG values when both α1 chains were mutated.

### Location of *GABRA2* mutations are equivalent to pathogenic variants in other Cys-loop receptors

We investigated if the seven mutations identified in GABA_A_ α2 had equivalent positions with pathogenic variants in other Cys-loop receptors reported in the literature. Pathogenic variants were defined as likely pathogenic or pathogenic in ClinVar and Disease in Humsavar. Mutations at the equivalent positions or one flanking amino acid in all the Cys-loop receptor genes were considered, using as reference the alignment in Supp. Figure 3B. Six of the seven variants had at least one pathogenic mutation at the equivalent or flanking position in other Cys-loop receptor genes. Additional information for the variants is shown in Table 1.

**Table 1.**
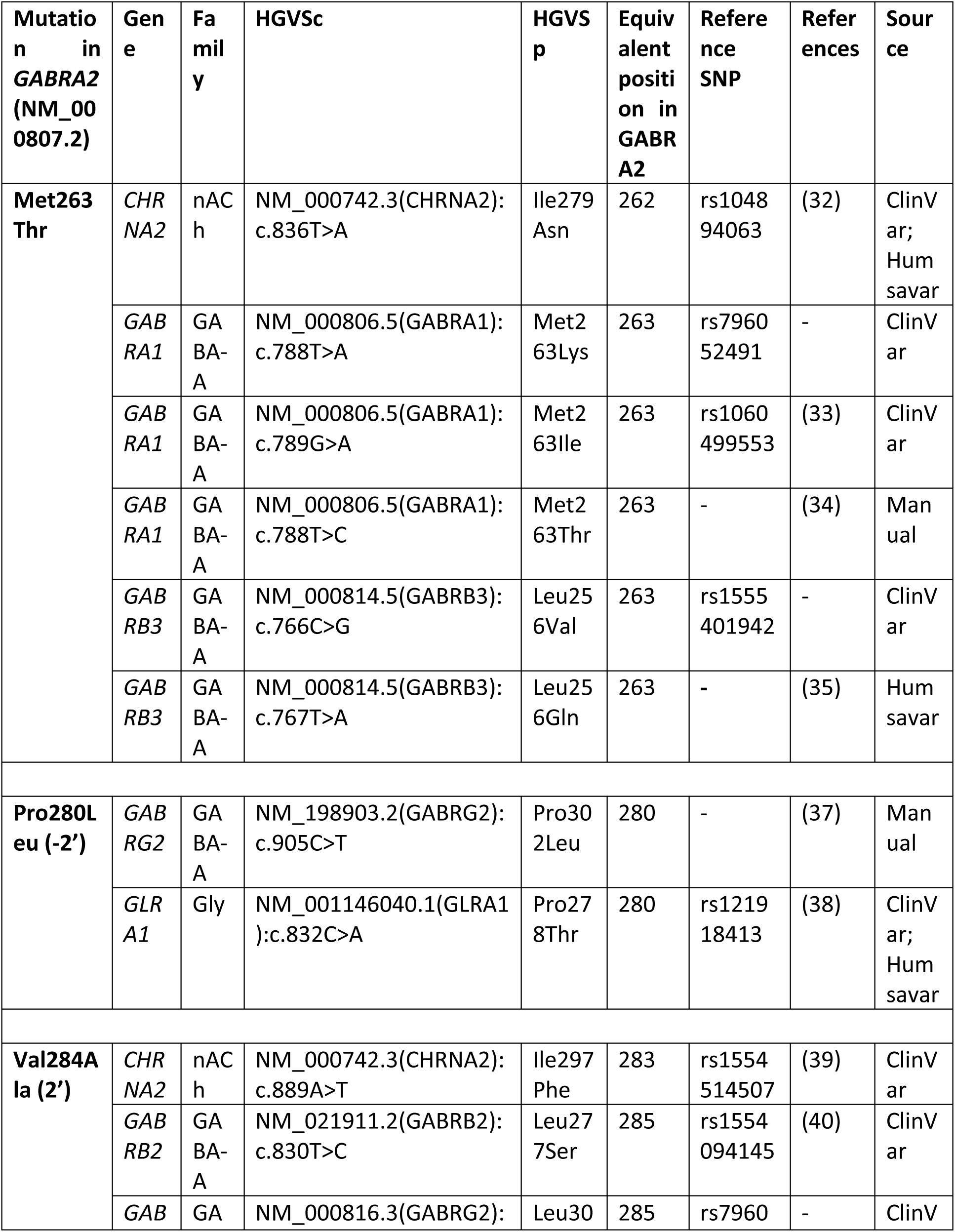
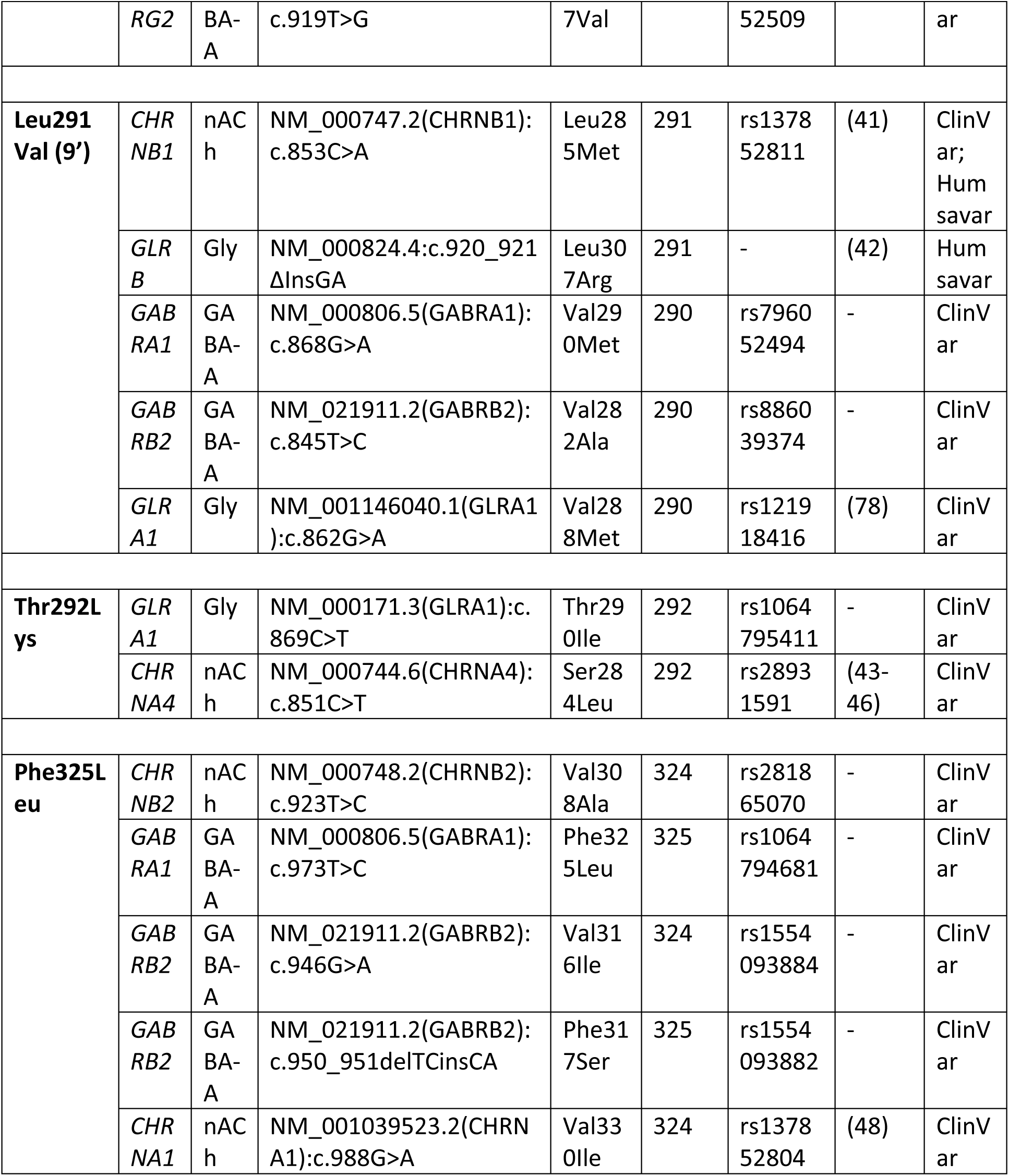
Details of variants in equivalent positions to the two previously reported and the novel mutation in *GABRA2* identified in other Cys-loop receptor genes.

For the Met263, five pathogenic mutations were observed to be equivalent to this exact position in other Cys-loop receptor genes, and one was present in the previous amino acid (32–35). This variant is in the M1 helix of the TM domain, in the inter-subunit interface with the M3, and this region is important for the actions of potentiating neuroactive steroids (36). Functional studies of the variant equivalent to the previous amino acid in *CHRNA2* (NM_000742.3:p.Ile279Asn), showed it increases the receptor sensitivity to the neurotransmitter, without affecting desensitization properties and channel permeability.

The same amino acid change than the novel variant in *GABRA2* (Pro280Leu) was reported at the equivalent location in *GABRG2* in an individual with Dravet Syndrome (37), and a different amino acid change was observed in an individual with autosomal dominant hyperekplexia (NM_001146040.1(*GLRA1*):p.Pro278Thr) (38). Both mutations, like the Pro280Leu, were in the −2’ ring of the receptors. The authors demonstrated that these mutations enhanced desensitization and reduced both the ‘channel open’ probability and the frequency of receptor single-channel openings.

Three variants reported in the literature were observed in the adjacent amino acids to the Val284Ala (2’) of *GABRA2* (Table 1) (39, 40). Functional studies on the NM_000742.3(*CHRNA2*):p.Ile297Phe were consistent with a LOF of the receptor, either by a possible impaired channel expression onto the cell surface or by a drastic decrease in the channel open probability (39).

Variants in the leucines that form the 9’ ring have been previously reported in other members of the Cys-loop receptors as associated with congenital myasthenic syndrome and Hyperekplexia (Table 1) (41, 42). Mutations at this site are known to destabilize the closed state of the receptor and produce spontaneously active channels. Functional works on the NM_000747.2(*CHRNB1*):p.Leu285Met (9’) showed reduced stability of the closed gate and pathologic channel openings even in the absence of the neurotransmitter, consistent with a receptor that is caught in the open state (41).

The equivalent position to Thr292 in GABA_A_ α2 has been reported as pathogenic in *GLRA1* (NM_000171.3:p.Thr290Ile) and *CHRNA4* (NM_000744.6:p.Ser284Leu) (43–46), being the last one a well-characterized variant identified in several unrelated families. Expression studies of the *CHRNA4*-Ser284Leu demonstrated higher affinity to acetylcholine and faster desensitization of the receptors (47).

Lastly, two variants were reported at the equivalent position to Phe325 in *GABRA2* and three were in the adjacent amino acid (Table 1). Functional studies on the variant in *CHRNA1* (NM_001039523.2:p.Val330Ile, equivalent to the adjacent Val324 in *GABRA2*) showed it affects the speed and efficiency of gating of its channel, slowing opening and increasing closing rates (48).

### Disease variants in TMH of Cys-loop receptors are most commonly found in M2

The previously reported and the novel variants in *GABRA2* fall in the TMH of the ion-channel. It was previously reported that TMHs are enriched for germline disease variants compared to other regions (49). Therefore, we investigated the TMH of all the Cys-loop receptors (Supp. Table 2) to contain pathogenic variants present in ClinVar and Humsavar.

The disease variants were more enriched in the TMH regions. Proteins were split into three features (i) non-TMH residues, (ii) TMH residues, and (iii) residues within ±5 residues of a TMH for which there were 55, 76, and 33 disease variants respectively. When normalizing the occurrence of the disease variants by the total number of residues present in the feature type across the Cys-loop receptors the non-TMH features had 0.0097 disease variants per residue, TMHs contained 0.058 disease variants per residue, and the ±5 flanking residues had 0.064 disease variants per residue. This establishes that disease variants were more than 5 times more likely to be a TMH residue than a non-TMH residue when accounting for the relative proportion of TMH residues of the total residues.

Within the TMH region, disease variants were more commonly found in, or in close proximity to, M2 than other TMH. 39 disease variants were observed in the M2 helix itself, and 47 when considering the ±5 flanking residues. M2 was the most populated helix for disease variants compared to M1, M3 and M4 (χ^2^ test p-value = 3.8e-7), and this effect was similar considering the TMH including the ±5 flanking residues (χ^2^ test p-value=7.3e-7) (Figure 4).

**Figure 4.**
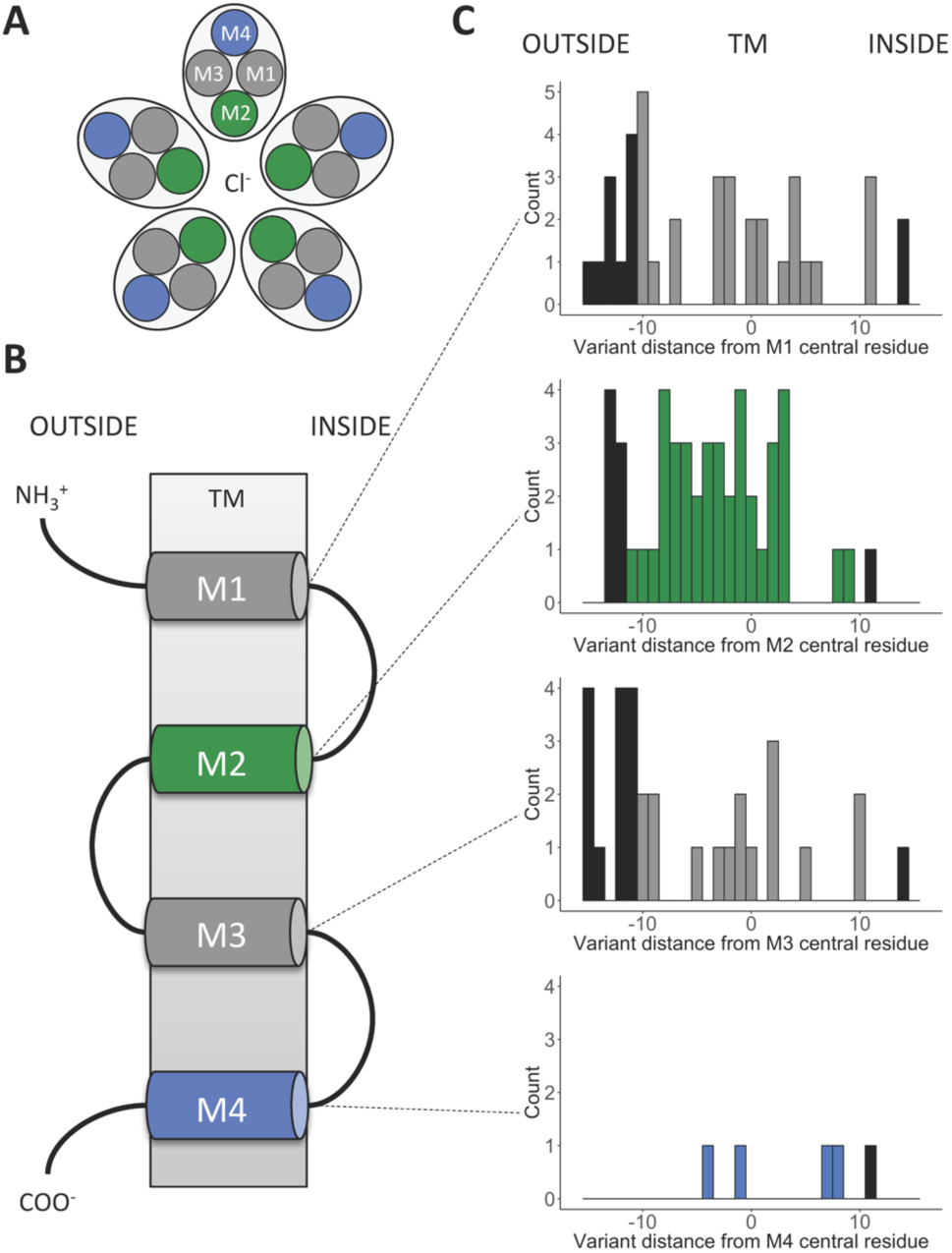
Distribution of disease variants along the TMH of Cys-loop receptors. **(A)** Top view schema of the TMH arrangement, considering the five subunits. **(B)** Simplified linear schema of the four helices of one subunit in the TM. **(C)** Disease variants positions plotted as a distance in residues from the central TM residue in sequence. Negative positions are towards the “outside” (extracellular space) whereas positive positions are towards the “inside” (cytosol). The number of variants observed in the TMH was 86: M1 = 27, M2 = 39, M3 = 16, M4 = 4, and 122 including ±5 flanking residues: M1 = 39, M2 = 47, M3 = 31, M4 = 5. In the histograms, disease variants in TMH are colored according to their position in the schema from Figure 4B, and disease variants in flanking regions are colored black. Duplicates in ClinVar and Humsavar are counted as one variant.

Notably, as a trend, the variants in M2 were distributed throughout the helix, whereas in M1 which is the second most populated helix for disease variants, these were most common at the lipid-water interface and flanking regions particularly on the extracellular side (a peak of 5 disease variants at position -10. Also, the M1 has 9 positions with no observed disease variants compared to 4 positions with no disease variants in the M2 (Figure 4C). M4 sustained fewer pathogenic variants, consistent with the lower conservation and higher tolerance for variation in gnomAD (Figure 1A, Supp. Figure 2).

## Discussion

Here we report a novel *de novo* missense mutation in *GABRA2* gene. The phenotype of this case was similar to the previously reported individuals, presenting with EIEE, developmental delay, significant hypotonia and congenital nystagmus. The variant (Pro280Leu) was present in the −2’ position of the channel pore (50). This position is important for the ion-selectivity and desensitization of the ion-channel, which is a fundamental property of most ligand-gated ion channels to change the conformation and limit the ion current flow, despite the neurotransmitter still being bound to the receptor (25). We observed that the variant was in a highly conserved residue and the radius of the pore was narrowed at the gate in both closed and desensitized structures of the receptor. Having a residue like proline with a rigid side chain (or an alanine like in the β subunits with a non-rigid but very small side chain) at the intracellular −2’ position allows tight control of the gating upon neurotransmitter extracellular binding, exerting rigid body motion of the TMH, as can be observed in recent GABA_A_ and Gly receptors structures (26–28, 51). Replacing the constraining proline with a leucine will permit changes in the backbone conformation (52). Moreover, ion-selectivity is determined by the −2’ Pro in anion-selective channels, along with the -1’ Ala (1, 53), with no requirement for a charged side chain in the narrowest constriction.

Mapping of the seven variants in *GABRA2* to the protein structure showed they are located near the desensitization gate, the activation gate, or the M1-M3 inter-subunit interfaces, which are key regions for the proper function of the receptor. Functional experiments of previously reported missense mutations in *GABRA2* have been seen to present both gain and loss of function properties, all associated with epilepsy. Variants reported by Maljevic *et al*. (21) were suggested to be compatible with a LOF mechanism, while the mutation in the activation gate (9’) reported by Butler *et al.* (20) was associated with predominantly opened channels, therefore, suggested to result in the gain-of-function (GOF) of the receptor. This could be explained by the dual role that *GABRA2* presents during development (54), since early in life GABA_A_ receptors mediate excitation by depolarizing immature neurons in response to activation by GABA, and this is changed when the mature chloride gradient is established later in life, when they mediate inhibition by exerting hyperpolarization.

Six of the seven variants identified in *GABRA2* had equivalent pathogenic mutations in other Cys-loop receptors reported in the literature (Table 1), highlighting the importance of these regions. Although distinct phenotypes associated with the variants in Cys-loop receptors reflect their unique biological role, this family share a common structural topology conserved throughout evolution. Here, we show how variant interpretation by gathering together sequence, evolutionary, and structural information from homologous Cys-loop receptors can facilitate the characterization of novel candidate missense variants, especially in those cases where pathogenic mutations have already been characterized.

Furthermore, analysis of the distribution of previously reported pathogenic variants in the Cys-loop receptors showed that mutations are more common in TMH domain, more specifically, in the M2 segment, consistent with the importance of this helix in shaping the pore along the TM region. Also, the distribution of variants in M2 occurs throughout the helix, while disease variants in other helices that are more distant from the axis of the pore favor the water-lipid interface regions rather than the center of the TMH. Other clusters of pathogenic variants were observed in the M1 and M3 helices. M1 and M3 are more distant from the axis of the pore but play an important role in defining the inter-subunit interaction interfaces (27). In contrast M4, which is less conserved and more tolerant to variation in gnomAD (Figure 1), has fewer pathogenic variants. These results were also consistent with the observed constraint for variation distribution in the protein structure (Supp. Figure 2), where the M2 and M3 are more constrained than M1 and M4. Although mutations in these helices have been studied and demonstrated to influence channel gating kinetics, agonist sensitivity, and create spontaneous opening channels in different members of the Cys-loop receptors (55, 56), their distribution in the protein structure across the different members of the family has never been assessed. Our results provide a better understanding of the structural location of pathogenic variants in the TMH of Cys-loop receptors, facilitating further interpretation of novel candidate disease-causing mutations.

Herein, we present a novel *de novo* missense variant in *GABRA2* in a patient with EIEE and perform protein structural analysis of the previously reported and the novel variants. Our results highlight the importance of performing structural analysis of *de novo* missense mutations in *GABRA2*, in order to provide a more accurate insight into the etiology of the disease, that might also lead to opportunities for personalized treatments.

## Materials and Methods

### Patient identification and consent

The proband and both unaffected parents were recruited to the NIHR BioResouce research study. All participants provided written informed consent to participate in the study. The study was approved by the East of England Cambridge South national institutional review board (13/EE/0325). The research conforms with the principles of the Declaration of Helsinki.

### Genomic analysis

WGS was performed on DNA extracted from whole blood at 30x coverage using Illumina HiSeq X Ten system (Illumina, Inc., San Diego, CA, USA) with 150bp paired-end reads. Reads were aligned to the human genome of reference GRCh37/hg19 using Isaac Aligner, and variants were called with both Isaac Variant Caller (57) and Platypus (http://github.com/andyrimmer/Platypus). Variant annotation was performed with Variant Effect Predictor (58), which included allelic population frequencies from gnomAD (22) and deleteriousness scores from CADD (59), Sift (60) and Polyphen-2 (61). Structural variants were also identified by Manta (62) and Canvas (63) algorithms, as described previously (64, 65). Trio analysis focused on *de novo* and rare biallelic variant discovery unrestricted by a gene list. Candidate variants were then confirmed by Sanger sequencing.

### Protein sequence conservation

Conservation of GABA_A_R α2 was analyzed across orthologue protein sequences for different model species and across paralogue Cys-loop receptors from human. For that, the canonical protein sequences were obtained from UniProtKB (66), then aligned using MAFFT (67) with default parameters. Alignments were visually inspected using JalView (68).

### GABA_A_R structural analysis

There is no experimentally determined structure for the human GABA_A_R α2 subunit, either in a homo- or hetero-pentameric complex. For the structural analysis of the previously reported and the novel variants we used the recently solved cryo-electron microscopy (cryo-EM) human hetero-pentameric GABA_A_R α1β3γ2 complexes in closed (PDB ID 6HUG, with resolution 3.1Å) and desensitized (PDB ID 6HUP and 6I53, with resolution 3.58Å and 3.2Å respectively) forms (28, 69). These structures were used because they represent the most abundant arrangement of adult GABA_A_R hetero-pentamers which is two α subunits, two β subunits and one γ subunit. Also, the previously reported and the novel variants were in the M1, M2 and M3 segments of the TM domain, and this region was observed, by paired local sequence alignment, to present 100% identical residues between human GABA_A_R α1 and GABA_A_R α2.

The closed form structure was solved in complex with the pore blocker picrotoxin and the Megabody Mb38 protein. The desensitized structures were captured in complex with Diazepam (DZP), GABA and Megabody Mb38 for the PDB ID 6HUP, and with Megabody Mb38 for the PDB ID 6I53.

Structural mapping and visual inspection of the variants in the cryo-EM GABA_A_R α1β3γ2 structure were performed using PyMOL (70). The electrostatic surface visualization was done using its APBS plugin (71, 72). Residue interactions were calculated using Residue Interaction Network Generator (RING) (73) and visualized also with PyMOL.

### Channel pore characterization

In order to observe the effect of the previously reported and novel variant in the pore shape through the ion channel of the GABA_A_R α1β3γ2 structures, we considered four different configurations, i) where there was no mutated α1 subunit, ii) where the mutated α1 subunit was between β3 and γ2 (α1^β3γ2^), iii) where the mutated α1 subunit was between β3 and β3 (α1^β3β3^), and iv) where both α1 subunits were mutated (α1^β3γ2-β3β3^). The three mutant structures for each closed or desensitized forms were generated with the *BuildModel* command of FoldX (74), with a previous minimization using *RepairPDB* of the same program. In both steps, the ‘*membrane’* parameter was turned on, and for *BuildModel* twenty runs were requested. Then, for each mutant configuration the structure with the lowest difference in free energy of unfolding (ΔΔG=ΔG_mutant_-ΔG_WT_) was selected and the radii along the pore axis were calculated in steps of 3Å with PoreWalker (75), and compared between them.

### Pathogenic variants in TMH of Cys-loop receptors

We characterized the distribution of pathogenic variants in other Cys-loop receptor proteins (Supp. Table 2). Pathogenic variants were obtained from ClinVar annotated as “Pathogenic”, “Likely pathogenic” or “Pathogenic/Likely pathogenic” (accessed 31-Jan-2019) (76), and Humsavar, annotated as “Disease”, (release 13-Feb-2019, https://www.uniprot.org/docs/humsavar.txt). Pathogenic variants were mapped to UniProt canonical sequences using VarMap (77). Only missense variants were considered. Duplicates between Humsavar and ClinVar were only counted once, where a duplicate was considered to be the same amino acid change at the same position in the same protein, and variants with conflicting interpretations between Humsavar and ClinVar were omitted. TRANSMEM annotation by SwissProt was used to determine TMH boundaries. 265 variants were within the TMH, of which 86 were annotated as disease-causing. 398 variants were found in the TMH including ±5 flanking residues, of which 122 were annotated as disease-causing, being 82 from ClinVar, 11 from Humsavar and 29 from both.

The χ^2^ test from the SciPy python package (https://docs.scipy.org/doc/scipy/reference/generated/scipy.stats.chisquare.html) was used to compare the observed frequencies of variants that caused disease in M1, M2, M3, and M4 with those expected by chance.

## Acknowledgements

We thank NIHR BioResource volunteers for their participation, and gratefully acknowledge NIHR BioResource centers, NHS Trusts and staff for their contribution. We thank the National Institute for Health Research and NHS Blood and Transplant. The views expressed are those of the author(s) and not necessarily those of the NHS, the NIHR or the Department of Health. We also thank Dr. Courtney E French for her valuable comments on the figures.

## Data Availability Statement

Sequence data for the trio have been deposited at the European Genome-Phenome Archive (EGAD00001004522).

## Author contributions

ASJ and FLR designed the study. ASJ performed sequencing data processing. ASJ and MAH performed the former analysis and investigation, under the supervision of FLR and JT. JAB performed the statistical analysis. AMT, KS, MK and STD recruited the family and collected the clinical data and sample. KB performed the Sanger sequencing confirmation of the presented variant. ASJ, MAH and JAB wrote the manuscript. All authors read and approved the final manuscript.

