## Supplementary Figures for "Structural analysis of pathogenic missense mutations in *GABRA2* and identification of a novel *de novo* variant in the desensitization gate"

**Corresponding author**

**Supplementary Figure 1. Sanger sequencing confirmation of the variant identified in *GABRA2*. A)** Family pedigree structure. **B)** Sanger sequencing confirmation results. The variant was observed to be *de novo* in the proband.

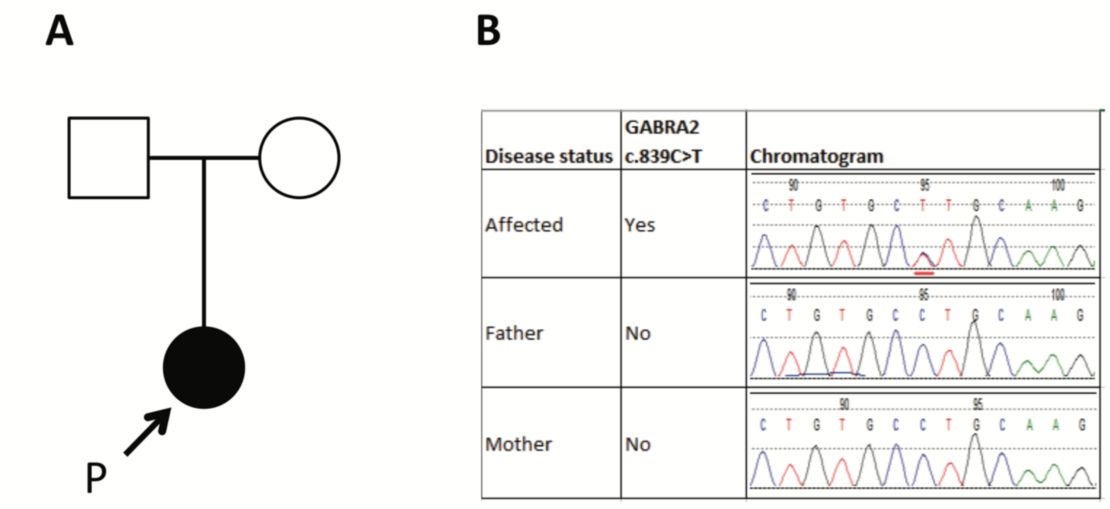

**Supplementary Figure 2. A) Constraint coding regions (CCRs) for *GABRA2* mapped in the GABA$_A$ α1β3γ2 receptor in desensitized state.** Colors for the CCRs match the ones used in Figure 1, and the variants highlighting is consistent with symbols as in Figure 1. The β3 chain is shown in blue and the γ2 is in yellow. **B)** Close view of one of the GABA binding sites. The GABA molecule is depicted in green. PDB ID of the GABA$_A$ α1β3γ2 receptor: 6HUP.

**A)** **B)**

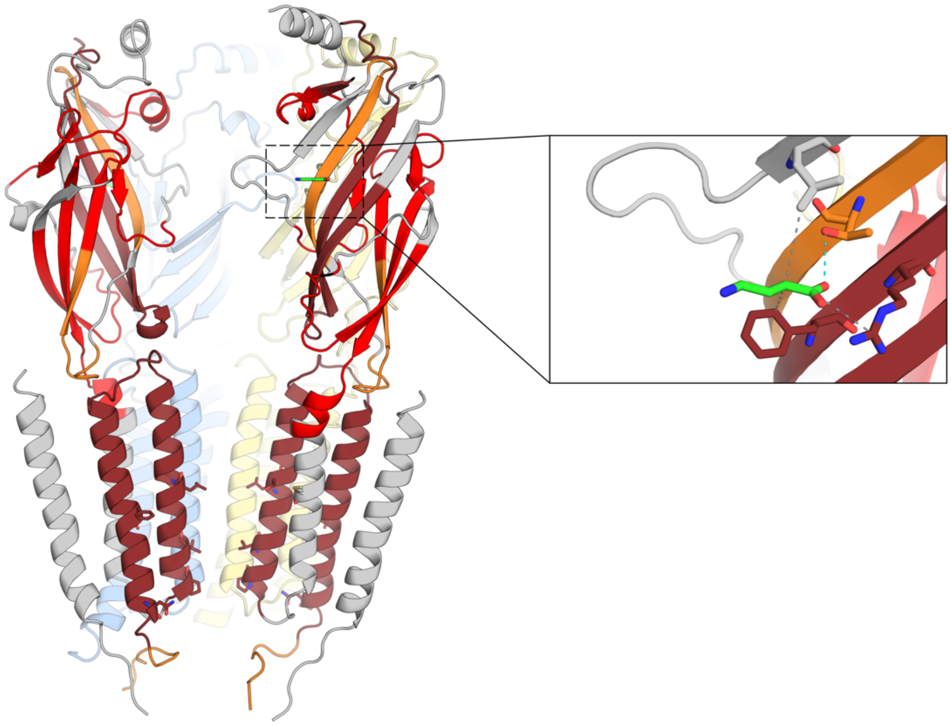

**Supplementary Figure 3. Multiple sequence alignment of the TM domain of GABA$_A$, Gly, nACh and 5-HT$_3$ members of the Cys-loop receptors from human.** The novel and previously reported variants in *GABRA2* are represented in the corresponding columns of the alignment, using the same symbols as in Figure 1. The approximate boundaries of the TMH, depicted with the violet bars, are based on GABA$_A$α1R structures (PDB IDs 6HUG, 6HUP and 6I53). **A)** Residues are colored according to their physicochemical properties (Jalview color scheme). **B)** Residues in in blue correspond with pathogenic variants in ClinVar and Humsavar. Residues in red correspond with the novel and previously reported variants in *GABRA2,* discussed in this study.

**A)**

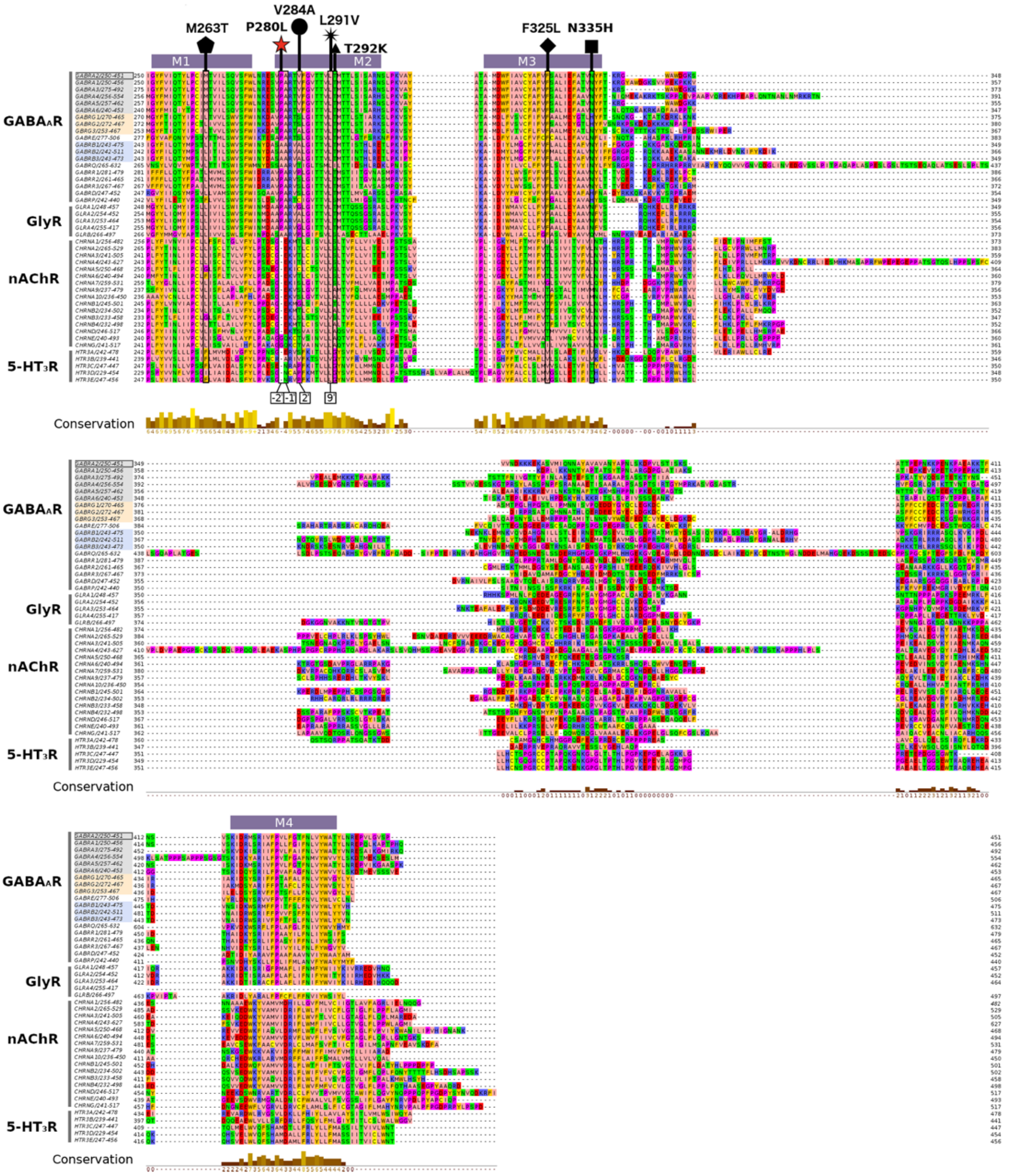

**B)**

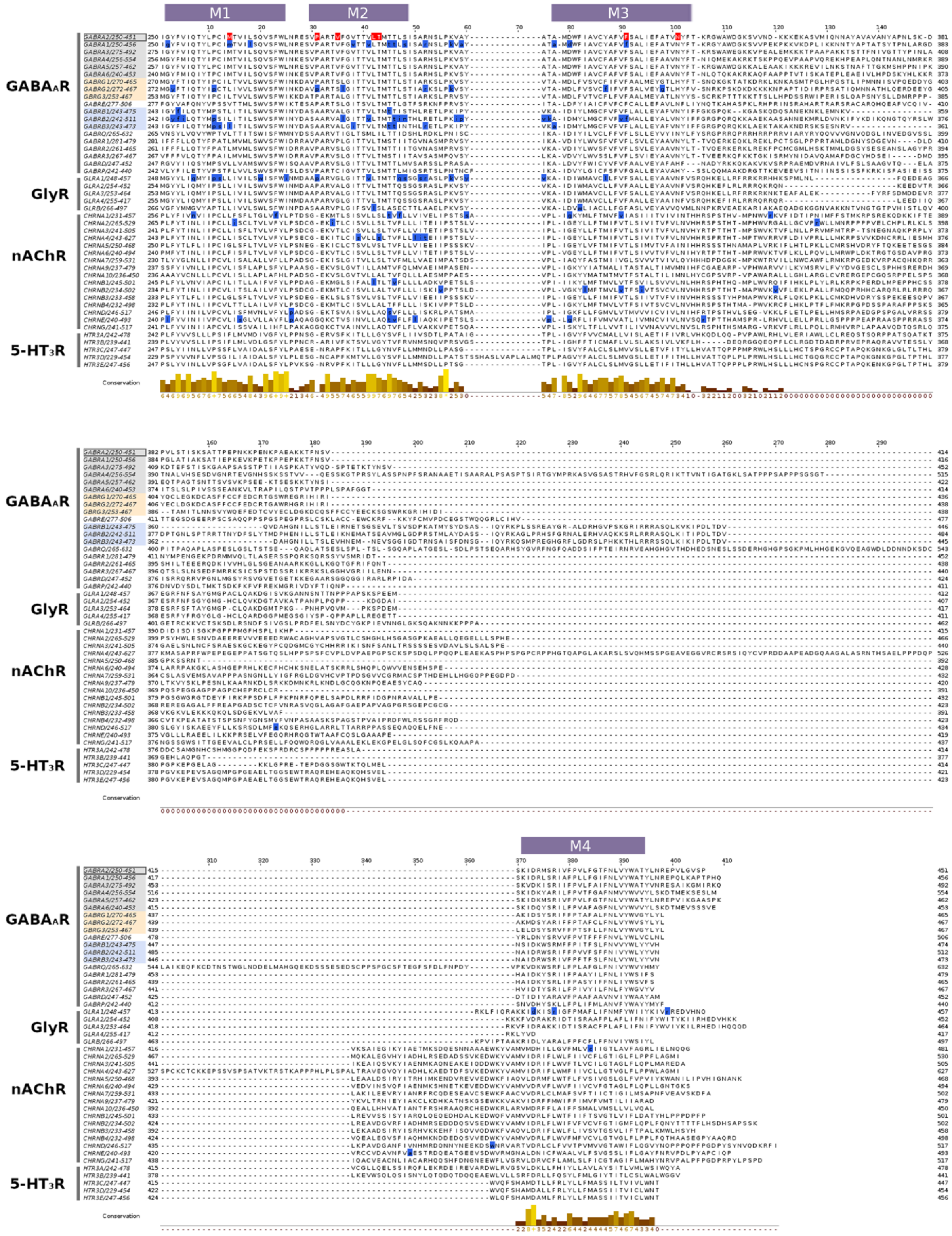

**Supplementary Figure 4. Radii profile measured at steps of 3 Å through the TM pore axis in the closed and desensitized states of the hetero-pentameric GABA$_A$ α1β3γ2 receptor.** The closed structure is in complex with PTX (PDB ID 6HUG). Two desensitized structures have been used, with and without GABA and DZP (PDB ID 6HUP and 6I53 respectively). **(A)** Profiles comparing the non-mutant structures. For the mutant structures, only the pore lining variants have been considered: **(B)** P280L, **(C)** V284A, **(D)** L291V, when the mutant α1 subunit is between β3γ2 (α1$^{β3γ2}$), between β3β3 (α1$^{β3β3}$), and when both α1 subunits are mutated (α1$^{β3γ2\text{-}β3β3}$). PTX=picrotoxin, DZP=diazepam.

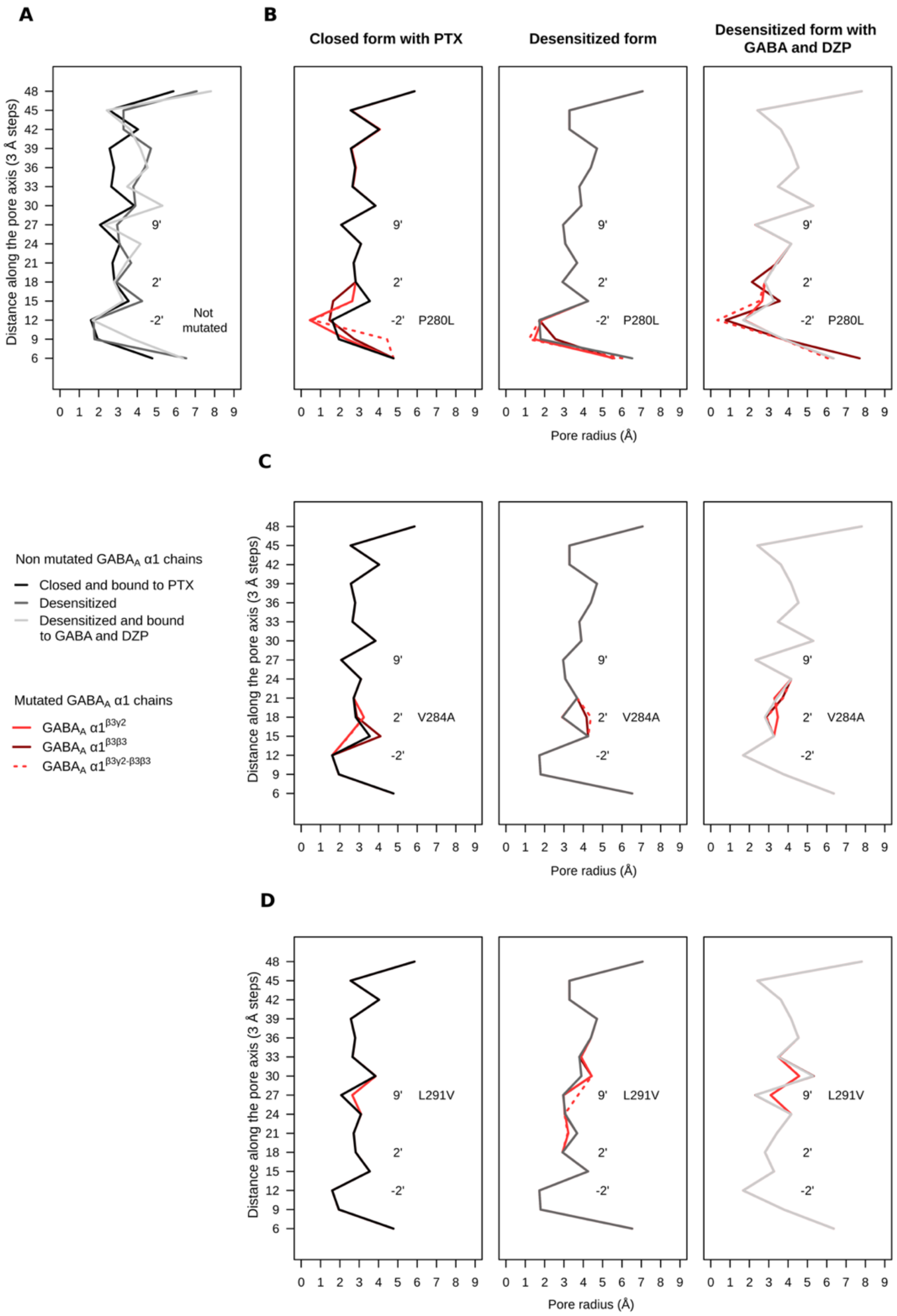

**Supplementary Figure 5. Anaesthetic-binding site of the diazepam (DZP) in the β3-α1 interface of the transmembrane (TM) domain of the $GABA_A$ α1β3γ2 receptor.** This receptor is in complex with DZP and GABA (PDB ID 6HUP). **A)** Side view of the β3-α1 interface. Exposed surface is colored according to electrostatic surface potential. **B)** Close view from the top of the cavity where the DZP (yellow) binds. DZP interacts directly with M263 and T292, and is close to the leucines that define the 9' ring. Symbols of the disease causing variants reported in *GABRA2* are the same as in Figure 1.

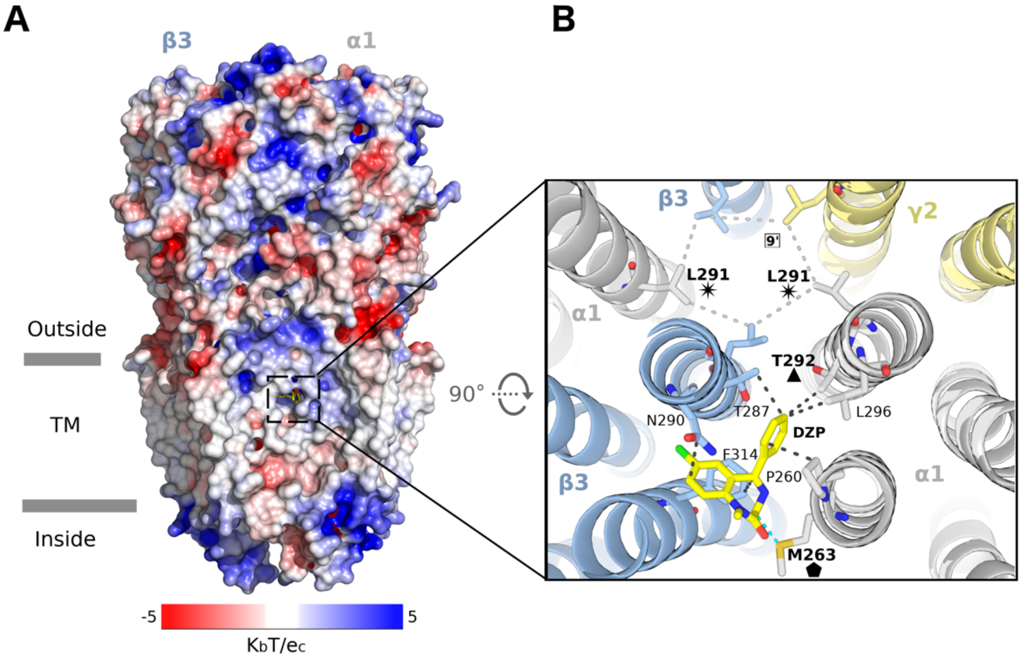
